# Lineage-Specific Silencing of *PSAT1* Induces Serine Auxotrophy and Sensitivity to Dietary Serine Starvation in Luminal Breast Tumors

**DOI:** 10.1101/2020.06.19.161844

**Authors:** Bo-Hyun Choi, Kelly O. Conger, Laura M. Selfors, Jonathan L. Coloff

## Abstract

A major challenge of targeting metabolism for cancer therapy is pathway redundancy, where multiple sources of critical nutrients can limit the effectiveness of some metabolism-targeted therapies. Here, we analyzed lineage-dependent gene expression in human breast tumors to identify differences in metabolic gene expression that may limit pathway redundancy and create therapeutic vulnerabilities. We found that the serine synthesis pathway gene *PSAT1* is the most depleted metabolic gene in luminal breast tumors relative to basal tumors. Low PSAT1 prevents *de novo* serine biosynthesis and sensitizes luminal breast cancer cells to serine and glycine starvation *in vitro* and *in vivo*. This *PSAT1* expression disparity—which pre-exists in the putative cells-of-origin of basal and luminal tumors—is due to luminal-specific hypermethylation of the *PSAT1* gene. Together, our data demonstrates that luminal breast tumors are auxotrophic for serine and may be uniquely sensitive to dietary serine starvation.

## INTRODUCTION

The unique metabolic phenotypes observed in cancer cells are driven by numerous factors, including genetic driver mutations, nutrient and oxygen availability, and the proliferation rate of the tumor (Vander Heiden and DeBerardinis, 2017). Recent work has also highlighted the extent to which tumor lineage (*i*.*e*., the cell- or tissue-of-origin) contributes to metabolic phenotypes in cancer (Mayers and Vander Heiden, 2017). This has been observed in human tumors at the gene expression level (Gaude and Frezza, 2016; Hu et al., 2013) as well as in controlled experiments demonstrating that the same driver mutations can induce distinct metabolic phenotypes depending on the tissue type in which they are activated (Mayers et al., 2016; Yuneva et al., 2012). The impact of tumor lineage on cancer phenotypes has also been emphasized by recent projects summarizing The Cancer Genome Atlas (TCGA) project, where a primary conclusion is that cell-of-origin gene expression patterns dominate the molecular classification of most tumors (Hoadley et al., 2018; Selfors et al., 2017).

Breast cancer is a highly heterogeneous disease made up of distinct subtypes of tumors. In the clinic, the histological expression of the estrogen receptor (ER), progesterone receptor (PR), and the receptor tyrosine kinase HER2 have guided therapeutic decisions for decades. Gene expression profiles have also been used to identify at least five molecular subtypes of breast cancer—basal, luminal A, luminal B, HER2+, and normal-like (Curtis et al., 2012; Lehmann et al., 2011; Perou et al., 2000; Sorlie et al., 2001). These molecular subtypes largely overlap with the clinical subtypes, with most luminal tumors being positive for ER and/or PR, and most basal tumors being triple negative (*i*.*e*., negative for ER, PR, and HER2) (Cancer Genome Atlas, 2012). More recently, molecular landscaping projects have simplified the classification of breast tumors into two primary lineages—luminal and basal—that are as distinct from each other as they are from tumors arising from completely distinct tissues-of-origin (Hoadley et al., 2018). When detected early, luminal tumors have a more favorable prognosis than basal tumors, in part due to their responsiveness to endocrine therapies. However, many luminal breast cancer patients suffer from relapse due to the development of therapeutic resistance (Osborne and Schiff, 2011). While the recent approvals of CDK4/6 and PI3K inhibitors have improved treatment of advanced luminal breast cancer, these therapies are still not curative for most patients (Andre et al., 2019; Finn et al., 2016; Goetz et al., 2017; Hortobagyi et al., 2018). As a result, luminal breast tumors still account for approximately half of all breast cancer fatalities (Siegel et al., 2019).

While recent research has identified many potential metabolic targets for cancer therapy, pathway redundancy has emerged as a major challenge of targeting metabolism for cancer therapy. The human genome, and in particular the human metabolic network, contains a high degree of functional redundancy (Nowak et al., 1997; Thiele et al., 2013). This plasticity is beneficial at the organismal level by allowing adaptation in a changing nutrient environment, but it complicates attempts to target metabolic pathways for cancer therapy. Here, we took the approach of analyzing metabolic gene expression in human breast tumors to identify cases where redundancy may be limited by lineage-dependent gene expression, thereby creating a metabolic vulnerability. Using this approach, we have found that the most significant differences in metabolic gene expression between luminal and basal breast tumors are found in the serine synthesis pathway, especially phosphoserine aminotransferase 1 (*PSAT1*), which is expressed at far lower levels in luminal tumors than basal tumors. Serine is a non-essential amino acid that is important for cancer cell proliferation not only for protein synthesis, but also for the synthesis of other amino acids, nucleotides, lipids, and antioxidants (Mattaini et al., 2016). Serine can either be taken up from the circulation or synthesized *de novo* via a three-step process catalyzed by phosphoglycerate dehydrogenase (PHGDH), PSAT1, and phosphoserine phosphatase (PSPH). Importantly, *PHGDH* amplifications and an enhanced ability to synthesize serine have previously been reported in basal breast tumors (Locasale et al., 2011; Possemato et al., 2011). This and other work motivated the development of PHGDH inhibitors as potential cancer treatments (Mullarky et al., 2016; Pacold et al., 2016; Rohde et al., 2018; Wang et al., 2017). While inhibition of serine synthesis has shown efficacy in some models and is still being evaluated, it is becoming clear that in some circumstances it is not effective due to extracellular serine that can be taken up to offset inhibition of *de novo* biosynthesis (Chen et al., 2013; Méndez-Lucas et al., 2020; Nilsson et al., 2012; Sullivan et al., 2019). Because our goal was to identify cases where lineage-dependent gene expression reduces pathway redundancy, we examined whether the very low expression of *PSAT1* found in luminal tumors limits their ability to synthesize serine and creates a dependency on exogenous serine for growth. We have found that luminal breast cancer cells are auxotrophic for serine due to lineage-specific hypermethylation of the *PSAT1* gene and are sensitive to serine starvation both *in vitro* and *in vivo*.

## RESULTS

### Lineage-specific suppression of the serine synthesis pathway in luminal breast cancer

To evaluate the relationship between tumor lineage and metabolic gene expression, we analyzed the expression of a curated list of 1454 metabolic genes (Gaude and Frezza, 2016) in The Cancer Genome Atlas (TCGA) Pan-Cancer Atlas data set (Hoadley et al., 2018). Unsupervised hierarchical clustering of metabolic genes in the 24 largest tumor types revealed that metabolic gene expression alone was largely sufficient to distinguish tumor tissue-of-origin (Figure 1A). Analyzing metabolic gene expression in breast tumors alone showed a clear distinction of the basal subtype, while luminal A and luminal B tumors largely clustered together (Figure S1A). Because luminal A and B tumors were similar with respect to metabolic gene expression, we focused our further analyses on luminal (*i*.*e*., luminal A + luminal B) and basal breast tumors, which cluster distinctly in our pan-cancer analysis of metabolic gene expression (Figure 1A).

**Figure 1.**
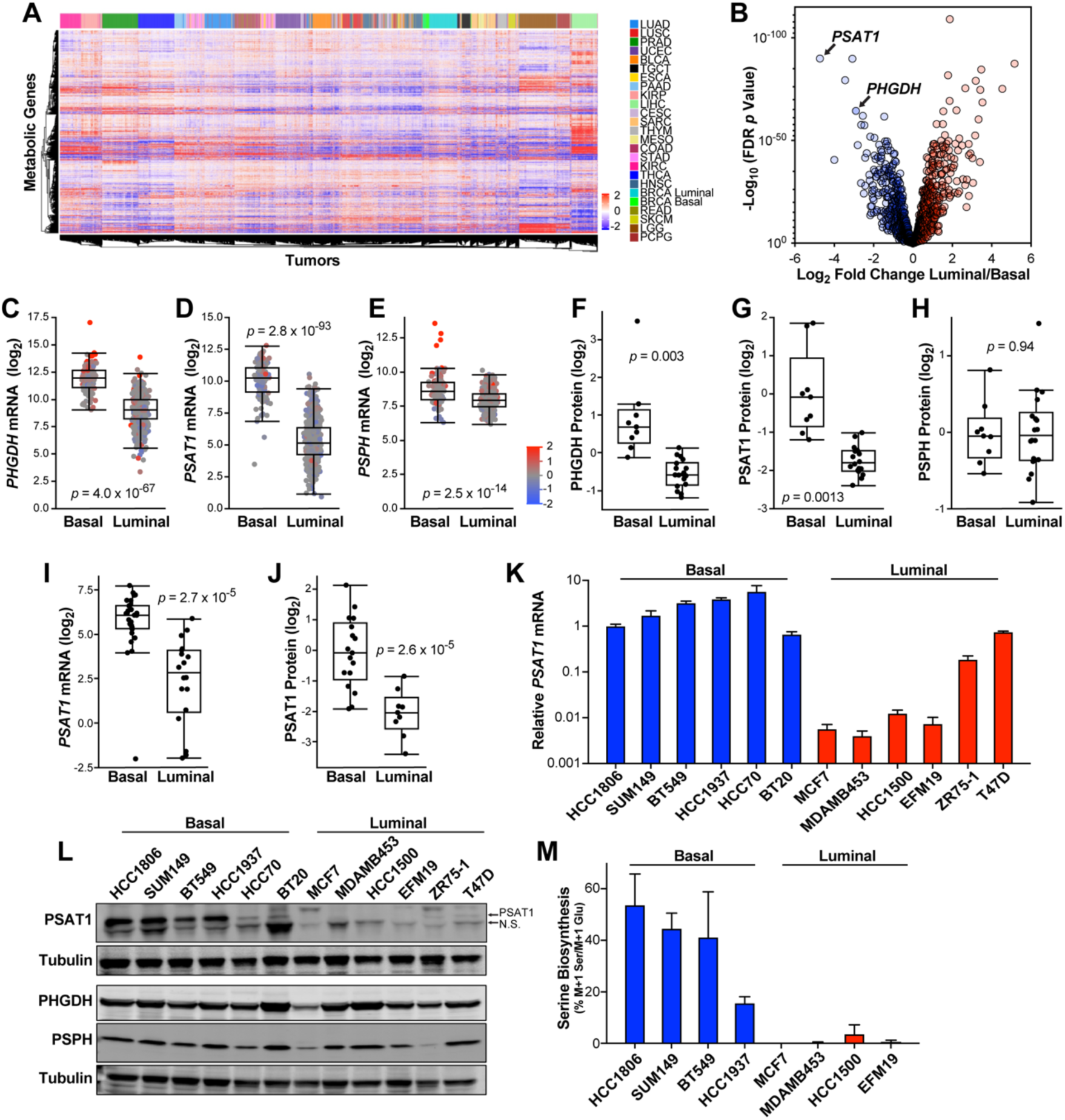
Lineage-Specific Suppression of the Serine Synthesis Pathway in Luminal Breast Tumors. **(A)** Hierarchical clustering of human tumors from the TCGA Pan-Cancer Atlas data set by metabolic gene expression. Data are log_2_ median centered. TCGA study abbreviations: LUAD = lung adenocarcinoma, LUSC = lung squamous cell carcinoma, PRAD = prostate adenocarcinoma, UCEC = uterine corpus endometrial carcinoma, BLCA = bladder urothelial carcinoma, TGCT = testicular germ cell tumors, ESCA = esophageal carcinoma, PAAD = pancreatic adenocarcinoma, KIRP = kidney renal papillary cell carcinoma, LIHC = liver hepatocellular carcinoma, CESC = cervical squamous cell carcinoma and endocervical adenocarcinoma, SARC = sarcoma, THYM = thymoma, MESO = mesothelioma, COAD = colon adenocarcinoma, STAD = stomach adenocarcinoma, KIRC = kidney renal clear cell carcinoma, THCA = thyroid carcinoma, HNSC = head and neck squamous cell carcinoma, BRCA = breast invasive carcinoma, READ = rectum adenocarcinoma, SKCM = skin cutaneous melanoma, LGG = brain lower grade glioma, PCPG = pheochromocytoma and paraganglioma. **(B)** Differences in metabolic gene expression between luminal and basal breast tumors in the TCGA Pan-Cancer Atlas RNAseq data set. Data is the log_2_ fold change of mean gene expression in luminal breast tumors relative to basal breast tumors. -Log_10_ *p* values from two-sided Welch’s t tests that have been corrected for false discovery using the Benjamani-Hochberg method. **(C – E)** mRNA levels of *PHGDH* **(C)**, *PSAT1* **(D)** and *PSPH* **(E)** in basal and luminal breast tumors in TCGA data. *p* values from two-sided Welch’s t tests. Tumors are colored based on gene copy number data. **(F – H)** Protein levels of PHGDH **(F)**, PSAT1 **(G)** and PSPH **(H)** in basal and luminal breast tumors in proteome analyses. *p* values from two-sided Welch’s t tests. **(I & J)** mRNA **(I)** and protein **(J)** levels of PSAT1 in basal and luminal breast cancer cell lines from the Cancer Cell Line Encyclopedia (CCLE). *p* values from two-sided Welch’s t tests. **(K)** *PSAT1* mRNA level in basal and luminal breast cancer cell lines. Values are the means ± SEM of three independent experiments. **(L)** Representative western blot of PHGDH, PSAT1, and PSPH in basal and luminal breast cancer cell lines. N.S. indicates non-specific band. **(M)** Serine biosynthesis in basal and luminal breast cancer cell lines. Values are the mean ± SEM of two independent experiments.

To focus on potential vulnerabilities created by lineage-dependent gene expression, we identified the most significant gene expression differences between luminal and basal tumors (Figure 1B). This analysis found that *PSAT1* is the most depleted metabolic gene, with luminal tumors expressing 26-fold less *PSAT1* than basal tumors (Figures 1B and 1D). *PHGDH* was also among the top scoring genes, and *PSPH* expression was also reduced in luminal breast tumors (Figures 1B, 1C, 1E). Similar results were found in another large human breast tumor data set (Figure S1B) (Curtis et al., 2012). Notably, these differences are not solely due to copy number alterations, as most tumors display these phenotypes in the absence of amplifications or deletions (Figures 1C–1E). Analysis of the proteome of human breast tumors (Johansson et al., 2019) also revealed very low levels of PHGDH and PSAT1 (but not PSPH) protein in luminal breast tumors (Figure 1F–1H). Using data from the Cancer Cell Line Encyclopedia (CCLE) (Barretina et al., 2012) and our own qPCR and western blots, we found that breast cancer cell lines robustly maintain the low-*PSAT1* phenotype of luminal tumors, while the differences in *PHGDH* and *PSPH* were less pronounced or absent (Figures 1I–1L, S1C–S1H). To determine if these gene expression differences result in relevant metabolic changes, we determined the percentage of intracellular serine that is synthesized in the serine synthesis pathway by tracing the incorporation of ^15^N from α-^15^N-glutamine into serine. Since nitrogen from glutamate is used by PSAT1 to generate serine, normalization of M+1 serine to M+1 glutamate describes an accurate fraction of intracellular serine made in the serine synthesis pathway. Consistent with our expression analyses, we detected high serine synthesis pathway activity in basal, but not luminal cell lines (Figure 1M).

### Luminal breast cancer cells are auxotrophic for serine

We next sought to determine whether luminal and basal breast cancer cells differ in their response to serine starvation, which inhibits the growth of some cancer cells *in vitro* and *in vivo* (Maddocks et al., 2013; Maddocks et al., 2017). Glycine is typically removed in these experiments because some tissues can convert glycine back to serine *in vivo*, even though glycine cannot replace serine to support cancer cell proliferation (Labuschagne et al., 2014). Because the non-physiological nutrient levels found in traditional tissue culture medium can impact cellular dependence on certain amino acids (Muir et al., 2017; Tardito et al., 2015), we first determined whether physiological medium (human plasma-like medium, or HPLM (Cantor et al., 2017)) impacts the response to serine and glycine (S/G) starvation. While breast cancer cells proliferate slightly faster in RPMI than HPLM in the presence of S/G, upon S/G starvation cells proliferate much more slowly in RPMI than HPLM (Figures S2A–S2D), indicating that the use of physiological medium allows for enhanced growth without S/G. While at present we do not fully understand the cause of these differences, we utilized the more physiologically relevant HPLM for the remainder of our experiments.

We performed S/G starvation on six basal (HCC1806, SUM149, BT549, HCC1937, HCC70, and BT20) and six luminal cancer cell lines (MCF7, MDAMB453, ZR75-1, EFM19, HCC1500, and T47D), where we found that all cell lines grow more slowly in the absence of S/G (Figures 2A-2C, S2E, S2F). Importantly, however, all basal cell lines were able to proliferate without S/G, while only one luminal cell line (T47D) showed this capability (Figures 2B, 2C). Interestingly, T47D cells have relatively high *PSAT1* expression that is more comparable to basal cell lines than other luminal lines (Figures 1K, 1L). S/G starvation increased serine synthesis pathway gene expression in most cell lines, but PSAT1 mRNA and protein remained very low in most luminal lines (Figures 2D, 2E, S2G, S2H). Moreover, long-term (30 day) S/G deprivation did not further increase *PSAT1* expression, indicating that the low *PSAT1* phenotype of luminal cells is durable under long-term selective pressure (Figure S2I). We also found that luminal breast cancer cells show a dramatic reduction in intracellular serine upon S/G starvation, while basal cells are able to maintain a much larger serine pool (Figure 2F). Consistently, ^15^N tracing revealed that most of the serine found in basal cells cultured without S/G was made in the serine synthesis pathway, and that *de novo* serine biosynthesis remained virtually undetectable in luminal cells grown in the absence of S/G (Figure 2F). These results indicate that most luminal breast cancer cell lines are auxotrophic for serine and are highly dependent on exogenous serine to support proliferation.

**Figure 2.**
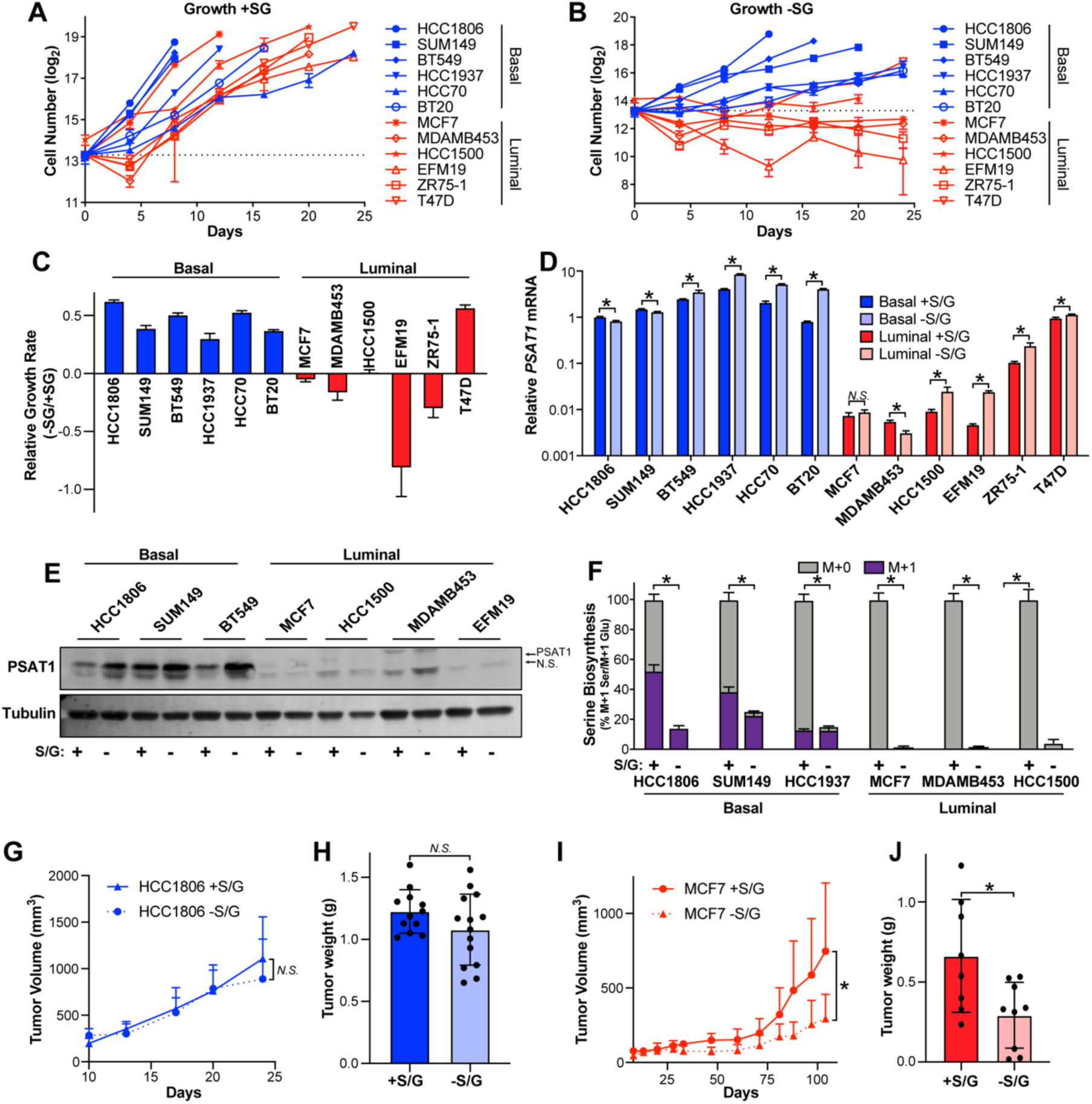
Luminal Breast Cancer Cells are Auxotrophic for Serine. **(A & B)** Basal and luminal breast cancer cell lines were cultured in normal HPLM media (+S/G) **(A)** or HPLM without S/G (-S/G) **(B)** and cell numbers were counted for 24 days or until cells became confluent. S/G starvation began on day 2. Dashed line indicates the number of cells plated per well. **(C)** Relative growth rates of basal and luminal breast cancer cells grown in the presence and absence of S/G. **(D)** *PSAT1* mRNA level in basal and luminal lines treated +/- S/G for 48 hrs. Values are the mean ± SD of triplicate samples from an experiment representative of three independent experiments. * indicates *p* < 0.05 from unpaired two-sided t tests. *N*.*S*. (not significant) indicates *p* > 0.05. **(E)** Representative western blot of basal and luminal cell lines treated +/- S/G for 48 hrs. **(F)** Serine abundance and biosynthesis in basal and luminal lines treated +/- S/G for 48 hrs. Values are the mean ± SD of triplicate samples from an experiment representative of two independent experiments. * indicates *p* < 0.05 from unpaired two-sided t tests on serine abundance data. **(G)** Tumor volume over time after injecting HCC1806 cells into the mammary gland of nude mice fed +/- S/G diet. Values are the mean ± SD. n = 12 tumors for +S/G and n = 14 tumors for -S/G. *N*.*S*. indicates p > 0.05 in mixed-effects model analysis. **(H)** HCC1806 tumor weight as measured at endpoint. Values are the mean ± SD (n = 12 tumors for +S/G and n = 14 tumors for -S/G). *N*.*S*. indicates p > 0.05 in two-sided Welch’s t test. **(I)** Tumor volume over time after injecting MCF7 cells into the mammary gland of the nude mice fed +/- S/G diet. Values are the mean ± SD (n = 8 tumors for +S/G and n = 9 tumors for -S/G). * indicates p < 0.05 in mixed-effects model analysis. **(J)** MCF7 tumor weight as measured at the endpoint. Values are the mean ± SD (n = 8 tumors for +S/G and n = 9 tumors for -S/G). * indicates p < 0.05 in two-sided Welch’s t test.

To determine whether the difference in basal and luminal cancer cell dependence on exogenous serine is recapitulated in the more complex *in vivo* environment, we injected basal HCC1806 and luminal MCF7 cells orthotopically into the mammary fat pads of nude mice and fed them custom diets with and without S/G. Consistent with previous reports, we found that a S/G free diet reduces plasma serine and glycine levels by ∼50% (Figures S2J, S2K). In line with our *in vitro* findings, the S/G-free diet had no effect on basal HCC1806 tumor growth but significantly inhibited the growth of luminal MCF7 tumors (Figure 2G–2J).

### PSAT1 expression determines growth in the absence of exogenous serine

Previous studies have demonstrated that several genetic features of tumors influence the response to serine starvation (DeNicola et al., 2015; LeBoeuf et al., 2020; Maddocks et al., 2013; Maddocks et al., 2017). In addition, the ability to synthesize serine *de novo* can allow cancer cells to proliferate when exogenous serine is limiting (Diehl et al., 2019; Méndez-Lucas et al., 2020; Sullivan et al., 2019). We examined whether the ability of basal breast cancer cells to proliferate without exogenous S/G is reliant on serine biosynthesis by treating basal HCC1806 cells with the PHGDH inhibitor PH-755 while growing +/- S/G. Despite strongly inhibiting serine synthesis, PH-755 had no effect on growth when S/G are present, but nearly completely blocked proliferation when extracellular S/G were absent (Figures S3A, S3B). Similarly, CRISPR-mediated knock out of either PHGDH or PSAT1 in HCC1806 and SUM149 cells had virtually no effect on proliferation in the presence of S/G, but completely halted growth upon S/G starvation (Figures 3A,3B, 3D, 3E). Accordingly, knockout of PHGDH or PSAT1 prevented serine synthesis and the maintenance of a serine pool in basal cells cultured without S/G (Figures 3C, 3F).

**Figure 3.**
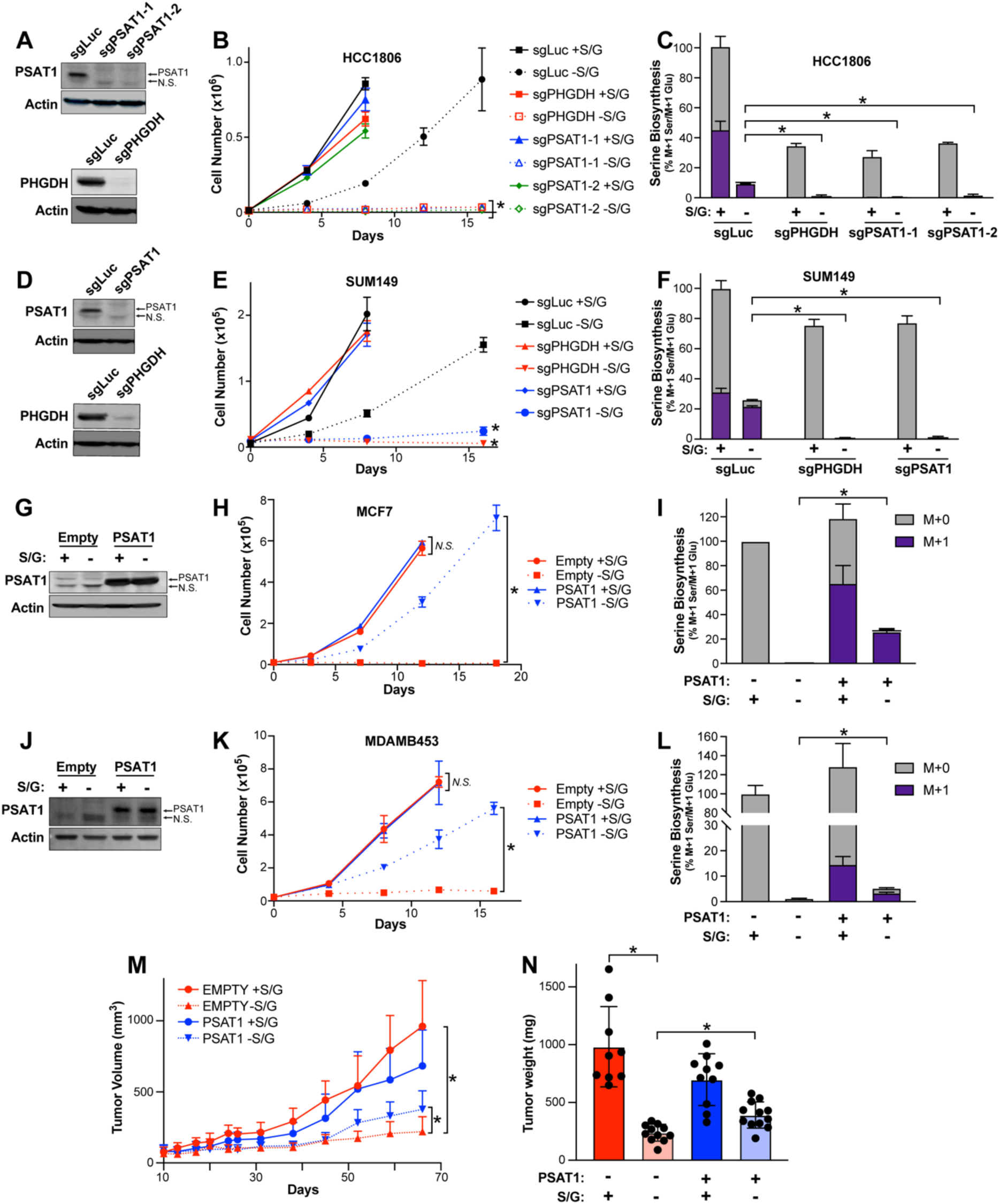
PSAT1 Expression Determines Growth in the Absence of Exogenous Serine. **(A)** Representative western blots of control (sgLuc), PSAT1 knockout (sgPSAT1-1 and sgPSAT1-2), and PHGDH knockout (sgPHGDH) HCC1806 cells. **(B)** Growth curve of control (sgLuc), sgPHGDH, and sgPSAT1 HCC1806 cells grown ± S/G. Values are the mean ± SD of triplicate samples from an experiment representative of three independent experiments. * indicates p < 0.05 in two-way repeated measures ANOVA. sgPSAT1 or sgPHGDH -S/G data were compared to sgLuc -S/G data. **(C)** Serine abundance and biosynthesis in sgLuc, sgPHGDH, and sgPSAT1 HCC1806 cells treated +/- S/G for 48 hrs. Values are the mean ± SD of triplicate samples from an experiment representative of two independent experiments. * indicates *p* < 0.05 from unpaired two-sided t tests on serine abundance data. **(D)** Representative western blots of control (sgLuc), PSAT1 knockout (sgPSAT1), and PHGDH knockout (sgPHGDH) SUM149 cells. **(E)** Growth curve of control (sgLuc), sgPHGDH, and sgPSAT1 SUM149 cells grown ± S/G. Values are the mean ± SD of triplicate samples from an experiment representative of three independent experiments. * indicates p < 0.05 in two-way repeated measures ANOVA. sgPSAT1 or sgPHGDH -S/G data were compared to sgLuc -S/G data. **(F)** Serine abundance and biosynthesis in sgLuc, sgPHGDH, and sgPSAT1 SUM149 cells treated +/- S/G f
or 48 hrs. Values are the mean ± SD of triplicate samples from an experiment representative of two independent experiments. * indicates *p* < 0.05 from unpaired two-sided t tests on serine abundance data. **(G)** Representative western blot of control (Empty) and PSAT1 overexpressing MCF7 (PSAT1) cells cultured ± S/G for 48 hrs. **(H)** Growth curve of Empty and PSAT1 MCF7 cells cultured ± S/G. Values are the mean ± SD of triplicate samples from one experiment representative of three independent experiments. * indicates *p* < 0.05 and *N*.*S*. indicates *p* > 0.05 in two-way repeated measures ANOVA tests comparing Empty and PSAT1 cells either + or – S/G. **(I)** Serine abundance and biosynthesis in Empty and PSAT1 MCF7 cells cultured +/- S/G for 48 hrs. Values are the mean ± SD of triplicate samples from an experiment representative of two independent experiments. * indicates *p* < 0.05 from unpaired two-sided t tests on serine abundance data. **(J)** Representative western blot of control (Empty) and PSAT1 overexpressing MDAMB453 cells (PSAT1) ± S/G for 48 hrs. **(K)** Growth curve of Empty and PSAT1 MDAMB453 cells cultured ± S/G. Values are the mean ± SD of triplicate samples from an experiment representative of three independent experiments. * indicates *p* < 0.05 and *N*.*S*. indicates *p* > 0.05 in two-way repeated measures ANOVA tests comparing Empty and PSAT1 cells either + or – S/G. **(L)** Serine abundance and biosynthesis in Empty and PSAT1 MDAMB453 cells cultured +/- S/G for 48 hrs. Values are the mean ± SD of triplicate samples from an experiment representative of two independent experiments. * indicates *p* < 0.05 from unpaired two-sided t tests on serine abundance data. **(M)** Tumor volume over time after injecting Empty and PSAT1 MCF7 cells into the mammary gland of the nude mice fed +/- S/G diet. Values are the mean ± SD (n = 9 tumors for Empty +S/G, n = 11 tumors for Empty -S/G, n = 10 tumors for PSAT1 +S/G and n = 12 tumors for PSAT1 -S/G). * indicates p < 0.05 in mixed-effects model analyses. **(N)** Empty and PSAT1 MCF7 xenograft tumor weight as measured at endpoint. Values are the mean ± SD (n = 9 tumors for Empty +S/G, n = 11 tumors for Empty -S/G, n = 10 tumors for PSAT1 +S/G and n = 12 tumors for PSAT1 -S/G). * indicates p < 0.05 in two-sided Welch’s t tests.

To determine whether the low PSAT1 expression of luminal cells contributes to their inability to grow in the absence of S/G, we overexpressed PSAT1 in the luminal cell lines MCF7 and MDAMB453 (Figures 3G, 3J). While not affecting growth +S/G, PSAT1 overexpression was sufficient to allow luminal cell proliferation without S/G (Figures 3H,3K). Furthermore, we found that PSAT1 overexpression was sufficient to allow *de novo* serine synthesis even in the presence of exogenous S/G and allowed for the maintenance of a relatively large serine pool upon S/G deprivation (Figures 3I, 3L). Treatment with PH-755 prevented PSAT1-induced growth -S/G, indicating that this rescue was specifically due to serine biosynthesis (Figure S3C). Importantly, PSAT1 overexpression was also sufficient to partially rescue the inhibition of MCF7 xenograft growth caused by dietary S/G starvation, demonstrating that at least part of the effect of S/G starvation is tumor cell intrinsic (Figures 3M, 3N). Together, these results indicate that PSAT1 expression is limiting for serine biosynthesis in luminal breast cancer cells and that low PSAT1 expression sensitizes luminal breast cancer cells to S/G deprivation *in vitro* and *in vivo*.

### *PSAT1* is epigenetically-silenced specifically in luminal breast tumors

Lineage-dependent gene expression is typically controlled by epigenetic modifications like DNA methylation (Kim and Costello, 2017). As such, we analyzed the TCGA human breast tumor DNA methylation data set, where we saw a strong correlation between the differences in metabolic gene expression and methylation between luminal and basal tumors (Figure 4A). Importantly, *PSAT1* is the most differentially methylated gene between luminal and basal tumors (Figure 4B). DNA methylation is strongly correlated with suppression of transcription, and high luminal *PSAT1* methylation is consistent with the low *PSAT1* seen in these tumors (Figures 1D, 4C) (Coloff et al., 2016). We also observed high *PSAT1* gene methylation specifically in luminal breast cancer cell lines using CCLE data and our own methylation-specific PCR assay (Figures 4D, 4E, S4A). Treatment of MCF7 cells with the DNA methyltransferase (DNMT) inhibitor azacytidine induced a strong induction of *PSAT1* mRNA despite only partially reducing *PSAT1* gene methylation, and was sufficient to induce serine biosynthesis (Figures 4F–4H). Treatment with another structurally distinct DNMT inhibitor also induced *PSAT1* mRNA expression in MCF7 cells (Figure S4B), and azacytidine and the related DNMT inhibitor decitabine also induced *PSAT1* expression in MDAMB453 cells (Figure S4C). These results demonstrate that *PSAT1* gene methylation contributes to the reduced *PSAT1* expression and low serine synthesis observed in luminal cells.

**Figure 4.**
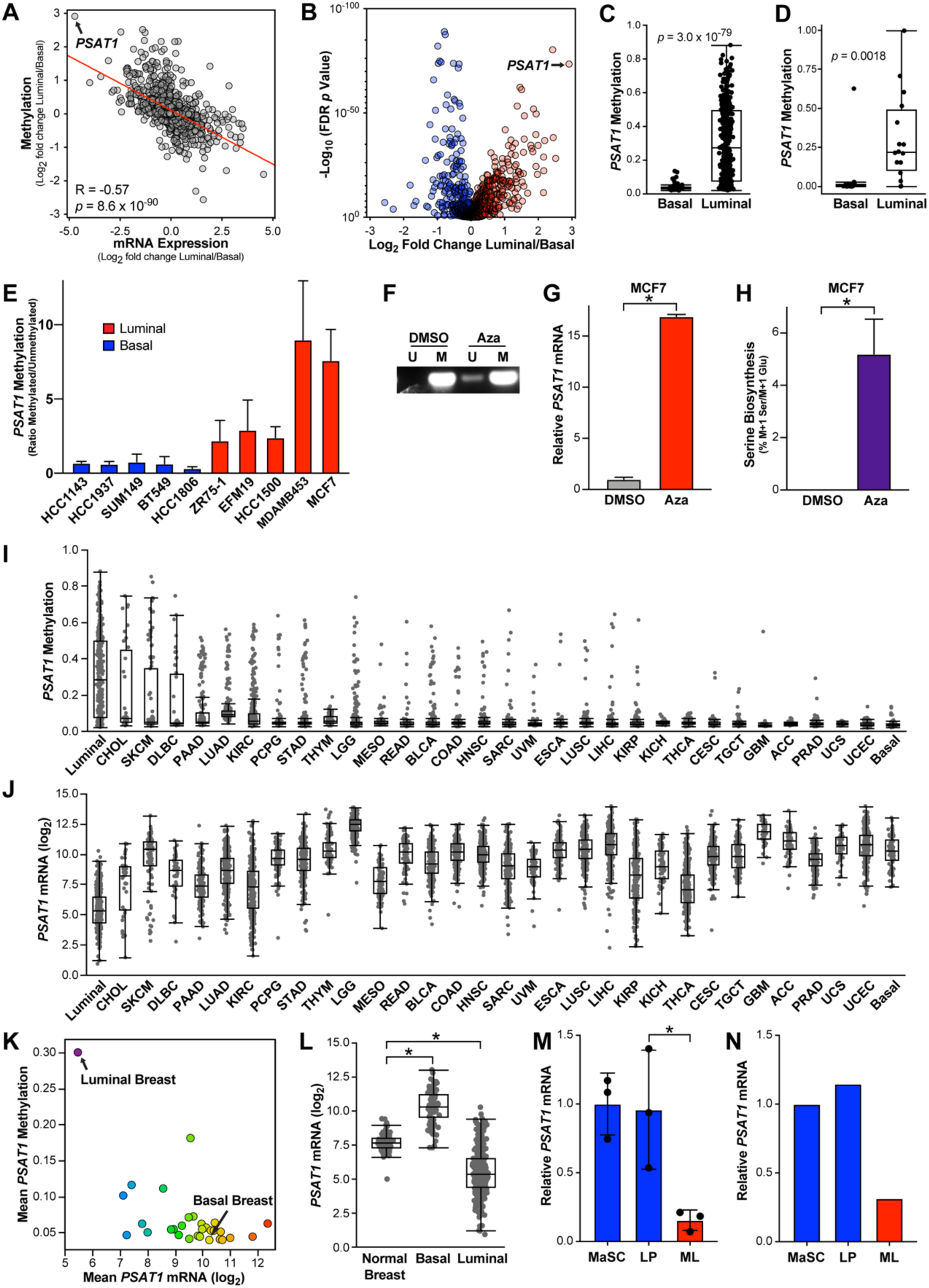
*PSAT1* is Epigenetically Silenced Specifically in Luminal Breast Cancer Cells. **(A)** Pearson correlation of fold changes in mRNA expression and DNA methylation between luminal and basal TCGA breast tumor samples. **(B)** Differences in metabolic gene methylation between luminal and basal breast tumors in the TCGA DNA methylation data set. Data is the log_2_ fold change of mean gene methylation in luminal breast tumors relative to basal breast tumors. -Log_10_ p values from two-sided Welch’s t tests that have been corrected for false discovery using the Benjamani-Hochberg method. **(C)** PSAT1 DNA methylation in basal and luminal breast tumors in the TCGA data set. *p* value from two-sided Welch’s t test. **(D)** *PSAT1* DNA methylation in basal and luminal breast cancer cell lines in the CCLE DNA methylation data set. *p* value from two-sided Welch’s t test. **(E)** Quantitation of the ratio of methylated over unmethylated *PSAT1* DNA in basal and luminal breast cancer cells. Values are the means ± SEM from two to four independent experiments. **(F)** Representative methylation specific PCR detecting methylated (M) and unmethylated (U) *PSAT1* promoter DNA in MCF7 treated with azacytidine 5 µM for 3 days. **(G)** *PSAT1* mRNA level in MCF7 cells treated with azacytidine (5 µM for 3 days). Values are the mean ± SD of triplicate samples from an experiment representative of three independent experiments. * indicates *p* < 0.05 in an unpaired two-sided t test. **(H)** Serine biosynthesis in MCF7 cells treated with azacytidine (5 µM for 3 days). Values are the mean ± SD of triplicate samples from an experiment representative of two independent experiments. * indicates *p* < 0.05 in an unpaired two-sided t test. **(I & J)** *PSAT1* gene methylation **(I)** and mRNA expression **(J)** by tumor type in the TCGA Pan-Cancer Atlas data sets. TCGA study abbreviations listed in legend for Figure 1A. **(K)** Mean PSAT1 methylation and mRNA expression by tumor type in the TCGA Pan-Cancer Atlas data sets. **(L)** *PSAT1* mRNA levels in normal breast tissue relative to basal and luminal breast tumors in TCGA breast cancer RNAseq data. * indicates *p* < 0.05 in two-sided Welch’s t tests. **(M)** Analysis of *PSAT1* mRNA expression in sorted luminal progenitor cell (LP) and mature luminal cells (ML) derived from normal human mammary glands. Microarray data from (Lim et al., 2009). * indicates *p* < 0.05 in an unpaired two-sided t test. **(N)** Analysis of *PSAT1* (J) mRNA expression in sorted LP and ML cells. RNAseq data from (Pellacani et al., 2016).

To determine whether high *PSAT1* methylation might induce sensitivity to S/G deprivation in additional tumor types, we analyzed *PSAT1* methylation and gene expression in the TCGA Pan-Cancer Atlas data set. Interestingly, luminal breast tumors have the highest *PSAT1* methylation and are the only tumor type to show consistently high *PSAT1* methylation, although outliers can be found in most tumor types (Figure 4I). Accordingly, luminal breast tumors have the lowest *PSAT1* mRNA expression of any tumor type (Figures 4J, 4K), suggesting that low *PSAT1*-induced sensitivity to S/G starvation may be a luminal breast tumor-specific vulnerability.

### The low *PSAT1* phenotype of luminal tumors pre-exists in their putative cells-of-origin

To understand how the low-*PSAT1* expression phenotype of luminal tumors arises, we investigated *PSAT1* expression in normal breast tissue. Interestingly, we found that normal breast expresses *PSAT1* at an intermediate level between that found in luminal and basal tumors (Figure 4L). This suggests that the low-*PSAT1* phenotype of luminal tumors either represents an active suppression of *PSAT1* or the outgrowth of a sub-population of mammary epithelial cells that express low levels of *PSAT1*. To investigate the latter possibility, we examined the expression of *PSAT1* in two independent data sets of FACS-sorted sub-populations of epithelial cells from normal breast reduction donors (Lim et al., 2009; Pellacani et al., 2016). The normal mammary gland is a complex structure that includes three major epithelial cell types—mammary stem cells, luminal progenitors, and mature luminal cells (Visvader and Stingl, 2014). It is believed that the cells-of-origin for basal tumors are similar to luminal progenitor cells and the cells-of-origin for luminal tumors are more similar to mature luminal cells (Figure S4D). In two independent data sets, we found that *PSAT1* expression is significantly higher in mammary stem cells and luminal progenitors than in mature luminal cells (Figures 4M, 4N). This data suggests that *PSAT1* expression is low in the cells-of-origin of luminal breast tumors, consistent with a model in which the serine auxotrophy of luminal tumors is driven by an outgrowth of luminal cells with a pre-existing low-*PSAT1* phenotype.

## DISCUSSION

Initial efforts of targeting serine metabolism in breast cancer were focused on “gain-of-function” cases where amplification of *PHGDH* leads to elevated rates of serine biosynthesis in basal tumors. However, it is becoming clear that targeting “gain-of-function” metabolic alterations in cancer can be challenging if redundant pathways remain present and active (Méndez-Lucas et al., 2020). Because our goal was to identify “loss-of-function” cases where pathway redundancy is limited by lineage-specific gene expression, we have identified a potentially more tractable opportunity to target serine metabolism in luminal, not basal, breast cancer.

Long-term serine starvation can lead to sensory defects in mice (Gantner et al., 2019), and it remains unclear whether altering dietary serine content alone can form the basis of an effective treatment in humans. Other approaches, like inhibition of serine transporters or enzymatic degradation of plasma serine might also be effective treatments based on the vulnerability we have identified. Interestingly, hypermethylation of the *PSAT1* gene in luminal breast tumors is reminiscent of T-cell acute lymphoblastic leukemia (T-ALL), where hypermethylation of the asparagine synthetase gene sensitizes T-ALL cells to degradation of plasma asparagine with L-asparaginase (Worton et al., 1991). In addition, it has recently been shown that serine availability may be uniquely low in the mammary gland relative to other tissues (Sullivan et al., 2019). A low serine environment combined with our discovery of serine auxotrophy in luminal breast tumors may provide a therapeutic window where serine metabolism can be effectively targeted without negative side-effects in luminal breast cancer.

In summary, our studies of lineage-dependent gene expression in breast cancer have revealed a novel vulnerability in serine metabolism specifically in luminal breast tumors. Lineage-specific suppression of *PSAT1* induces serine auxotrophy in luminal breast cancer cells and sensitizes them serine starvation. These findings demonstrate that lineage-dependent gene expression is sufficient to limit pathway redundancy and create therapeutic vulnerabilities that could be taken advantage of to target specific subtypes of tumors.

## ACKNOWLEDGMENTS

We would like to thank Vipin Suri, Nello Mainolfi, Adam Friedman, and Mark Manfredi from Raze Therapeutics, Inc., as well as Huiping Zhao, Debra Tonetti, Wen Gu, Bookyung Ko, Isaac Harris, and Christian Dibble for reagents, technical assistance, and/or helpful conversations. We also thank Joan Brugge and the Ludwig Center at Harvard for their support. This research was supported by the National Cancer Institute (1K22CA215828-01A1) (J.L.C.) and a Pilot Award from the University of Illinois at Chicago Breast Cancer Research Group (J.L.C.).

## AUTHOR CONTRIBUTIONS

Conceptualization: B.C., J.L.C.; Methodology, Formal Analysis, Investigation, Data Curation, Visualization, & Writing – Review & Editing: B.C., L.M.S., J.L.C.; Validation, B.C., K.O.C., J.L.C.; Writing – Original Draft, B.C., J.L.C.; Funding Acquisition, J.L.C.

## DECLARATION OF INTERESTS

None.

## STAR METHODS

### Mouse Procedures

All mouse procedures were approved by the University of Illinois at Chicago Animal Care Committee. HCC1806 (1×10^6^), MCF7 (3×10^6^), MCF7-EMPTY (5×10^6^) and MCF7-PSAT1 (5×10^6^) cells were injected into the fat pad of the #4 mammary glands of 6 to 8-week-old athymic nude-foxn1^nu^ mice (Envigo). Cell suspensions were injected in a volume of 50 μL growth factor-reduced Matrigel (Corning). Estrogen (E2) was administered via silastic capsules implanted subcutaneously as previously described (Molloy et al., 2014). Even though not required for HCC1806 tumor growth, E2 was provided for all experiments to maintain constant conditions. All tumors were monitored by caliper measurements over time and tumor volume was calculated using the formula ½ (width^2^ x length). Mice were euthanized according to institutional guidelines. Serine and glycine free (TD.160752) and control (TD.110839) diets were purchased from Envigo and diet formulations are listed as reported (Sullivan et al., 2019). Custom diets were administered five days after surgery and replaced at least weekly.

### Western Blots

Cells were lysed in mammalian cell lysis buffer (50mM Tris pH 7.5, 150mM NaCl and 0.5% NP40) containing a protease and phosphatase inhibitor cocktail (Bimake.com) and 1 µM MG132 (Selleckchem). Protein concentration was determined by BCA assay (Thermo Fisher). Quantified protein samples were separated by electrophoresis on 4−20% ready-made Tris-Glycine gels (Invitrogen) and transferred to PVDF membranes (Millipore). Membranes were blocked with 2% bovine serum albumin for 1 h and incubated overnight with one or more primary antibody: PSAT1 (Thermo Fisher, PA5-22124.), PHGDH (Sigma, HPA021241), PSPH (Santa Cruz, sc-365183), and Actin (Sigma, A1978). Overexpression and knockout of PSAT1 (Figure 3) confirmed the correct band on PSAT1 western blots. Membranes were washed with tween 20-containing tris buffered saline and incubated with fluorescence-conjugated secondary antibodies (Bio-Rad). Rhodamine-conjugated anti-tubulin was also treated as a secondary antibody (Bio-Rad, 12004166). Images were detected using a ChemiDoc MP Imaging System (Bio-Rad).

### RT-qPCR

RNA was isolated using Trizol reagent (Thermo Fisher) and cDNA was generated using qScript cDNA Synthesis Kits (Quantabio). RT-qPCR was performed with SYBR Green on an ABI ViiA7 real-time PCR system (Applied Biosystems), and results were normalized to the expression of *RPLPO*. The primer sequences were: *PSAT1-F: 5’-CGGTCCTGGAATACAAGGTG-3’; PSAT1-R: 5’-AACCAAGCCCATGACGTAGA-3’; PHGDH-F:5’-ATCTCTCACGGGGGTTGTG-3’; PHGDH-R: 5’-AGGCTCGCATCAGTGTCC-3’; PSPH-F: 5’-TGGAGATGGTGCCACAGATA-3’; PSPH-R: 5’-CCTCCAAATCCAATGAAAGC-3’ and RPLPO-F: 5’-ACGGGTACAAACGAGTCCTG-3’ and RPLPO-R: 5’-CGACTCTTCCTTGGCTTCAA-3’*.

### Cell Culture and Media

All cell lines were authenticated by STR analysis and were regularly tested for mycoplasma using the MycoAlert Mycoplasma Detection Kit (Lonza). Cells were grown in human plasma-like medium according to the published formulation (Cantor et al., 2017) with 5% dialyzed FBS (Sigma) and Pen/Strep (Invitrogen). Media was changed at least every two days. As needed, cells were incubated in RPMI media (R9010-01, US Biological Life Sciences) with or without serine and glycine with 5% dialyzed FBS. Cells were counted using a Z1 Coulter Particle Counter (Beckman Coulter) and growth rates were calculated using the following formula: growth rate = ln(final cell number/initial cell number)/time.

### GC-MS Metabolite Analyses

Cells were incubated in media containing α-^15^N-glutamine (Cambridge Isotope Laboratories) for 24 hours. Cells were lysed on ice in methanol, water, and chloroform. Norvaline was used as an internal standard. For mouse plasma analysis, plasma was mixed with isotopically labeled amino acid standards (Cambridge Isotopes) for absolute quantification. Metabolites were extracted by adding HPLC grade ethanol (Sigma-Aldrich), vortexing, and centrifuging at 21000 x g at 4°C for 10 min. Extracts were air dried and derivatized with MOX (Thermo Fisher, PI45950) and *N*-*tert*-butyldimethylsilyl-*N*-methyltrifluoroacetamide with 1% *tert*-butyldimethylchlorosilane (Sigma-Aldrich). Samples were analyzed by GC/MS using a HP-5MS Ultra Inert GC column (19091S-433UI, Agilent Technologies) installed in an Agilent 7890B gas chromatograph coupled to an Agilent 5977B mass spectrometer. Helium was used as the carrier gas. One microliter of sample was injected at 280°C. After injection, the GC oven was held at 60°C for 1 min. The oven was then ramped to 320°C at 10 °C/min. and held for 9 min. at 320°C. The MS system operated under electron impact ionization mode at 70 eV and the MS source and quadrupole were held at 230°C and 150°C respectively. Mass isotopomer distributions were determined by manually integrating ion fragments using ChemStation software (Agilent). Natural abundance correction was performed in R. Total abundance was normalized to the norvaline internal standard and to cell number counted in proxy wells. Serine biosynthesis was calculated by determining the fraction of labeled (M+1) serine and dividing it by the fraction of M+1 glutamate.

### Knockout and Overexpression

Knockout of PHGDH and PSAT1 was performed using lentiCRISPR v2 Puro (Addgene, 52961). The following oligos were cloned into BsmBI cut lentiCRISPR v2 sgLuc F: 5’-caccgGAGGCTAAGCGTCGCAA-3’; sgLuc R: 5’-aaacTTGCGACGCTTAGCCTCc-3’; sgPSAT1-1 F: 5’-caccgACCGAGGGGCACTCTCGG-3’; sgPSAT1-1 R: 5’-aaacCCGAGAGTGCCCCTCGGTc-3; sgPSAT1-2 F: 5’-caccgCATCACGGACAATCACCA-3’; sgPSAT1-2 R: 5’-aaacTGGTGATTGTCCGTGATGc-3’; sgPHGDH F: 5’-caccgAGTCTGGCCAGTGTGCCG-3’; sgPHGDH R: 5’-aaacCGGCACACTGGCCAGACTc-3’. For PSAT1 overexpression, human PSAT1 was cloned into pLenti CMV Neo DEST (Addgene, 17392) using the Gateway cloning system (Thermo Fisher). To generate lentiviral particles, HEK293T cells were transduced with PAX2, VSVG, and the lentiviral plasmid of interest using polyethylenimine (Polysciences). Viral supernatants were collected 48 and 72 hours after transduction. After infection with polybrene (hexadimethrine bromide, Sigma), cells were drug selected until mock infected cells were completely cleared after which antibiotic was removed.

### Methylation Specific PCR

Bisulfite conversion of genomic DNA was carried out using the EZ DNA Methylation-Gold kit (ZYMO Research Corporation) according to the manufacturer’s instructions. The following MSP primers were designed using MethPrimer 2.0 (Li and Dahiya, 2002): methylated *PSAT1* F 5’-GTAGGGTTTGCGATAGTACGG-3’; methylated *PSAT1* R 5’-GCTACGATAAAAATCTACAACCGAC-3’; unmethylated *PSAT1* F 5’-GGGTTTGTGATAGTATGGGT-3’; unmethylated *PSAT1* R 5’-CCACTACAATAAAAATCTACAACCAAC-3’. MSP PCR conditions consisted of a denaturing step of 15 min at 95°C followed by 40-50 cycles of 30s at 95°C, 30s at 59°C and 30s at 72°C, with a final extension of 7 min at 72°C. PCR products were analyzed by running on a 2% agarose gel with SYBR-Safe DNA gel stain (Invitrogen).

### Bioinformatics and Statistical Analyses

TCGA Pan-Cancer Atlas RNAseq (RSEM), DNA methylation, and copy number (gistic2) data sets were downloaded from the University of California, Santa Cruz Xena browser (xena.ucsc.edu). TCGA study abbreviations are: LUAD = lung adenocarcinoma, LUSC = lung squamous cell carcinoma, PRAD = prostate adenocarcinoma, UCEC = uterine corpus endometrial carcinoma, BLCA = bladder urothelial carcinoma, TGCT = testicular germ cell tumors, ESCA = esophageal carcinoma, PAAD = pancreatic adenocarcinoma, KIRP = kidney renal papillary cell carcinoma, LIHC = liver hepatocellular carcinoma, CESC = cervical squamous cell carcinoma and endocervical adenocarcinoma, SARC = sarcoma, THYM = thymoma, MESO = mesothelioma, COAD = colon adenocarcinoma, STAD = stomach adenocarcinoma, KIRC = kidney renal clear cell carcinoma, THCA = thyroid carcinoma, HNSC = head and neck squamous cell carcinoma, BRCA = breast invasive carcinoma, READ = rectum adenocarcinoma, SKCM = skin cutaneous melanoma, LGG = brain lower grade glioma, PCPG = pheochromocytoma and paraganglioma. CCLE RNAseq and DNA methylation data was downloaded from the CCLE web portal (portals.broadinstitute.org/ccle). Low detection genes, defined as >5% tumors with 0 counts, were removed from the analysis. METABRIC data was downloaded from cBioPortal (cbioportal.org). PAM50 subtype calls for breast tumors and cell lines were obtained from UCSC Xena and (Jiang et al., 2016), respectively. For basal vs luminal analyses, luminal A and B samples were grouped together and compared to basal samples, while HER2+ and normal-like samples were excluded. For each metabolic gene, the DNA methylation probe that most strongly anti-correlated with mRNA levels for that gene (lowest Pearson’s R) was selected. Hierarchical clustering and heatmap generation was performed using R 3.5.1. Data for heatmaps are log_2_ median centered. Box plots, volcano plots, and scatter plots were generated using JMP Pro 12 and GraphPad Prism 8. Statistical analyses were performed using JMP Pro 12, GraphPad Prism 8, and Microsoft Excel. Where applicable, the Benjamini-Hochberg procedure was used to correct for false discovery rate. Error bars represent either the SD or SEM as described in the figure legends. The Geiser-Greenhouse correction was used for two-way repeated measures ANOVA and linear mixed model analyses.

**Supplementary Figure 1.**
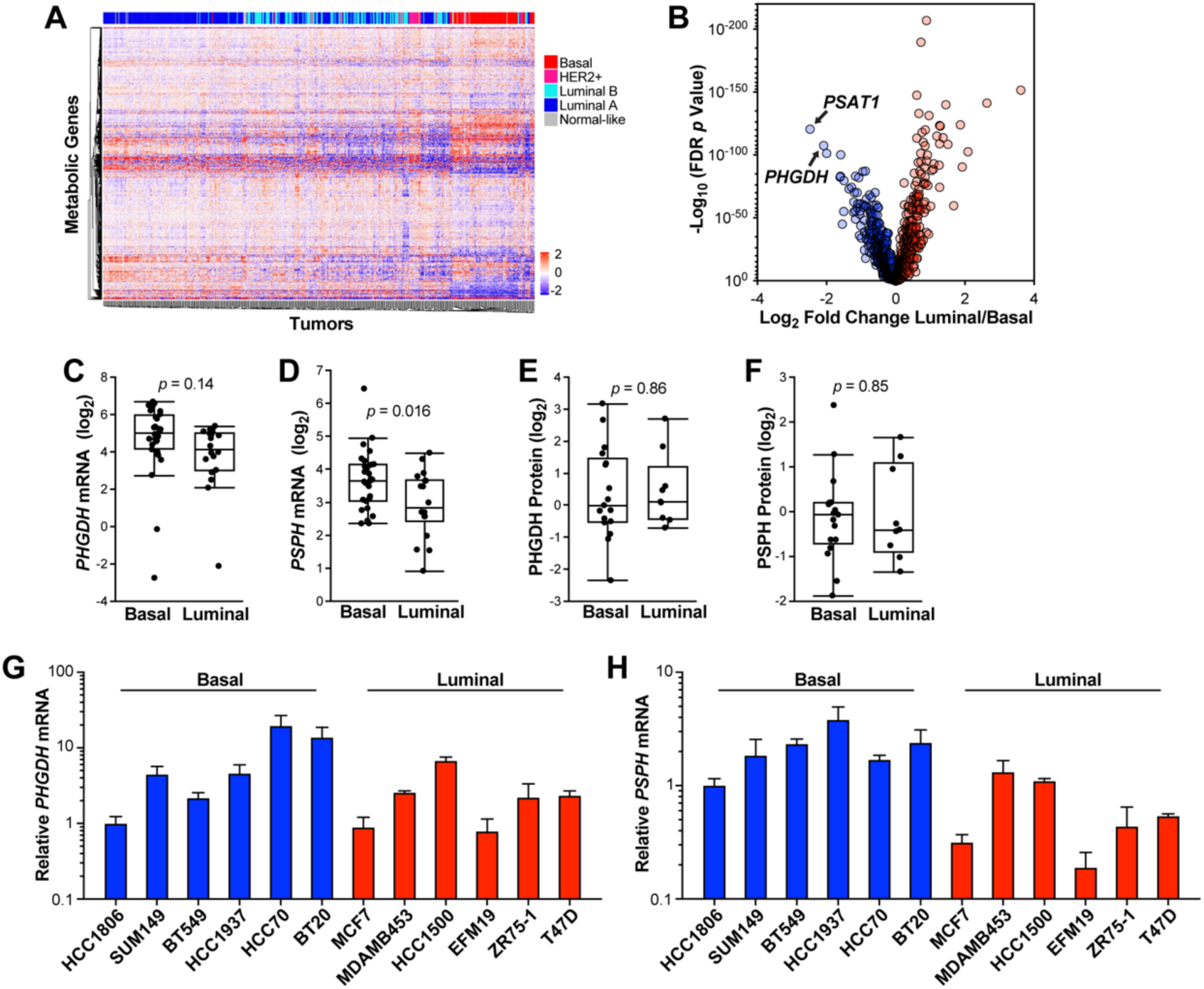
**Related to Figure 1.** **(A)** Hierarchical clustering of human breast tumors from the TCGA Pan-Cancer Atlas data set by metabolic gene expression. Data are log2-median centered. **(B)** Differences in metabolic gene expression between luminal and basal breast tumors in the METABRIC data set. Data is the log2 fold change of mean gene expression in luminal breast tumors relative to basal breast tumors. -Log10 p values from two-sided Welch’s t tests that have been corrected for false discovery using the Benjamani-Hochberg method. **(C & D)** *PHGDH* **(C)** and *PSPH* **(D)** mRNA levels in basal and luminal breast cancer cell lines from the Cancer Cell Line Encyclopedia (CCLE). Values are the means ± SEM of three independent experiments. *p* values from two-sided Welch’s t tests. **(E & F)** PHGDH **(E)** and PSPH **(F)** protein levels in basal and luminal breast cancer cell lines from the Cancer Cell Line Encyclopedia (CCLE). *p* values from two-sided Welch’s t tests. **(G & H)** *PHGDH* **(G)** and *PSPH* **(H)** mRNA levels in basal and luminal breast cancer cell lines. Values are the means ± SEM of three independent experiments.

**Supplementary Figure 2.**
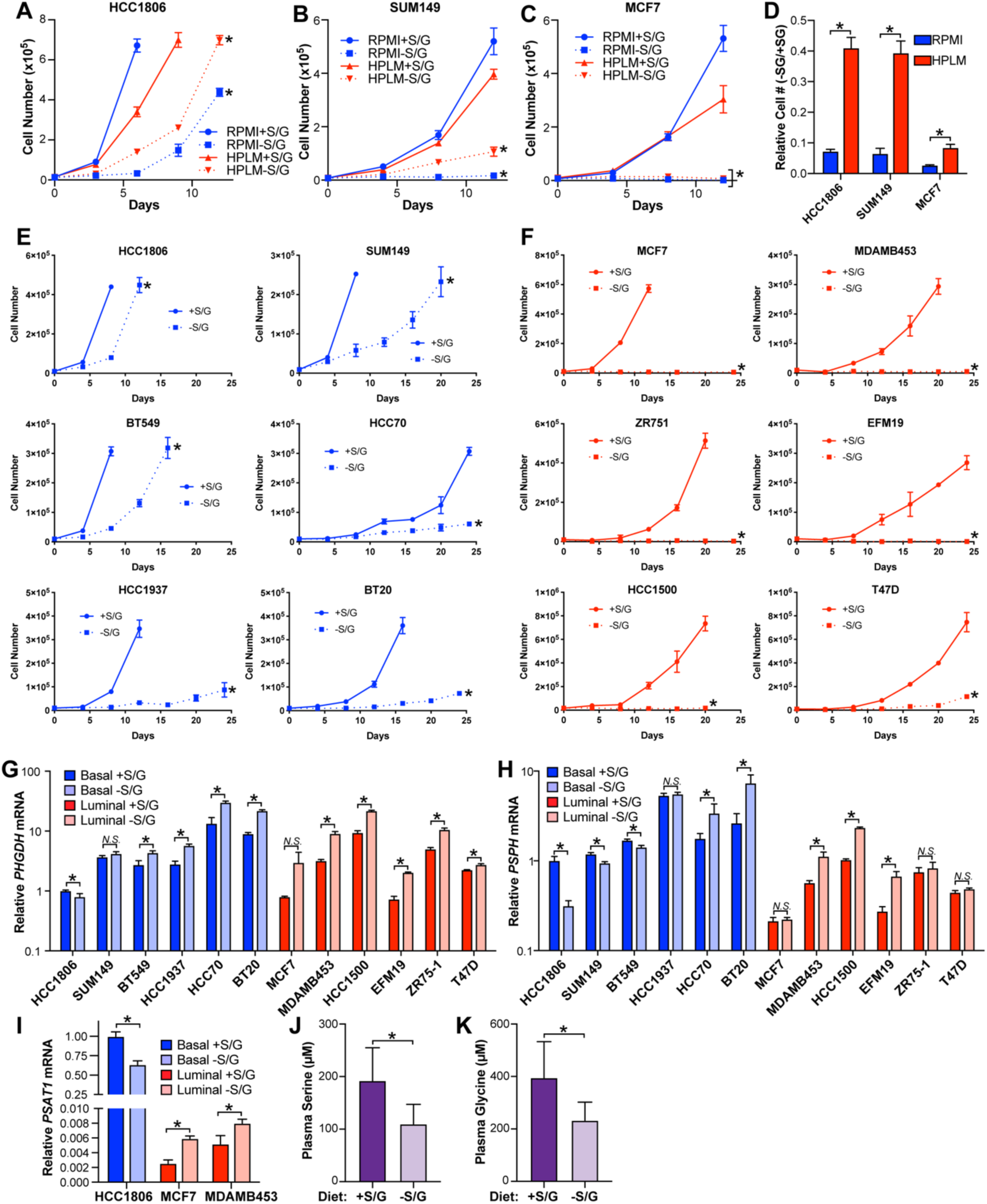
**Related to Figure 2.** **(A – C)** Growth curves of HCC1806 **(A)**, SUM149 **(B)**, and MCF7 **(C)** cultured in RPMI or HPLM in the presence or absence of S/G. * indicates *p* < 0.05 in two-way repeated measures ANOVA tests comparing +S/G to –S/G samples. **(D)** Ratio of cell numbers after 8 days of culture -S/G relative to +S/G in RPMI (blue) or HPLM (red). * indicates *p* < 0.05 in an unpaired two-sided t test. **(E & F)** Growth curves of basal **(E)** and luminal **(F)** breast cancer cells cultured in HPLM media ±S/G. Values are the means ± SD of one experiment representative of three independent experiments. * indicates *p* < 0.05 in two-way repeated measures ANOVA tests. **(G & H)** *PHGDH* **(G)** and *PSPH* **(H)** mRNA levels in basal and luminal lines treated +/- S/G for 48 hrs. Values are the mean ± SD of triplicate samples from an experiment representative of three independent experiments. * indicates *p* < 0.05 from unpaired two-sided t tests. *N*.*S*. (not significant) indicates *p* > 0.05. **(I)** *PSAT1* mRNA levels in breast cancer cells incubated +/- S/G for 30 days. Values are the mean +/- SD of triplicate samples from an experiment representative of two independent experiments. **(J & K)** Serine **(J)** and glycine **(K)** concentrations in mouse plasma after 21 days on custom +/- SG diets. n = 5 mice. * indicates *p* < 0.05 in an unpaired two-sided t test.

**Supplementary Figure 3.**
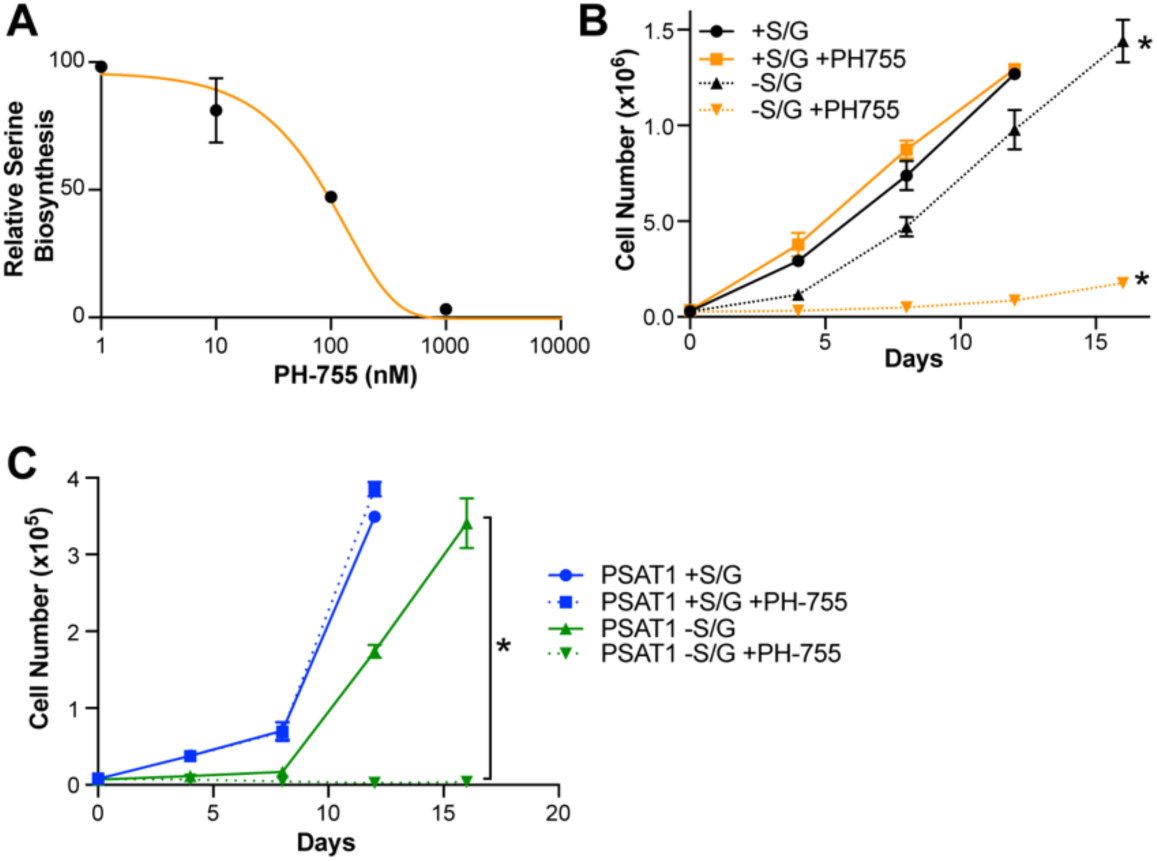
**Related to Figure 3.** **(A)** Relative labeled serine levels after culture with PH-755 at the indicated doses. Values are the mean ± SD of triplicate samples from an experiment representative of two independent experiments. **(B)** Proliferation of HCC1806 cells treated +/- 1 µM PH-755 +/- S/G. Values are the mean ± SD of triplicate samples from an experiment representative of three independent experiments. * indicates p <0.05 in two-way repeated measures ANOVA. -S/G data is compared to corresponding +S/G data. **(C)** Growth curve of PSAT1 overexpressing MCF7 cells treated +/- 1 µM PH-755 and S/G. Values are the mean ± SD of triplicate samples.

**Supplementary Figure 4.**
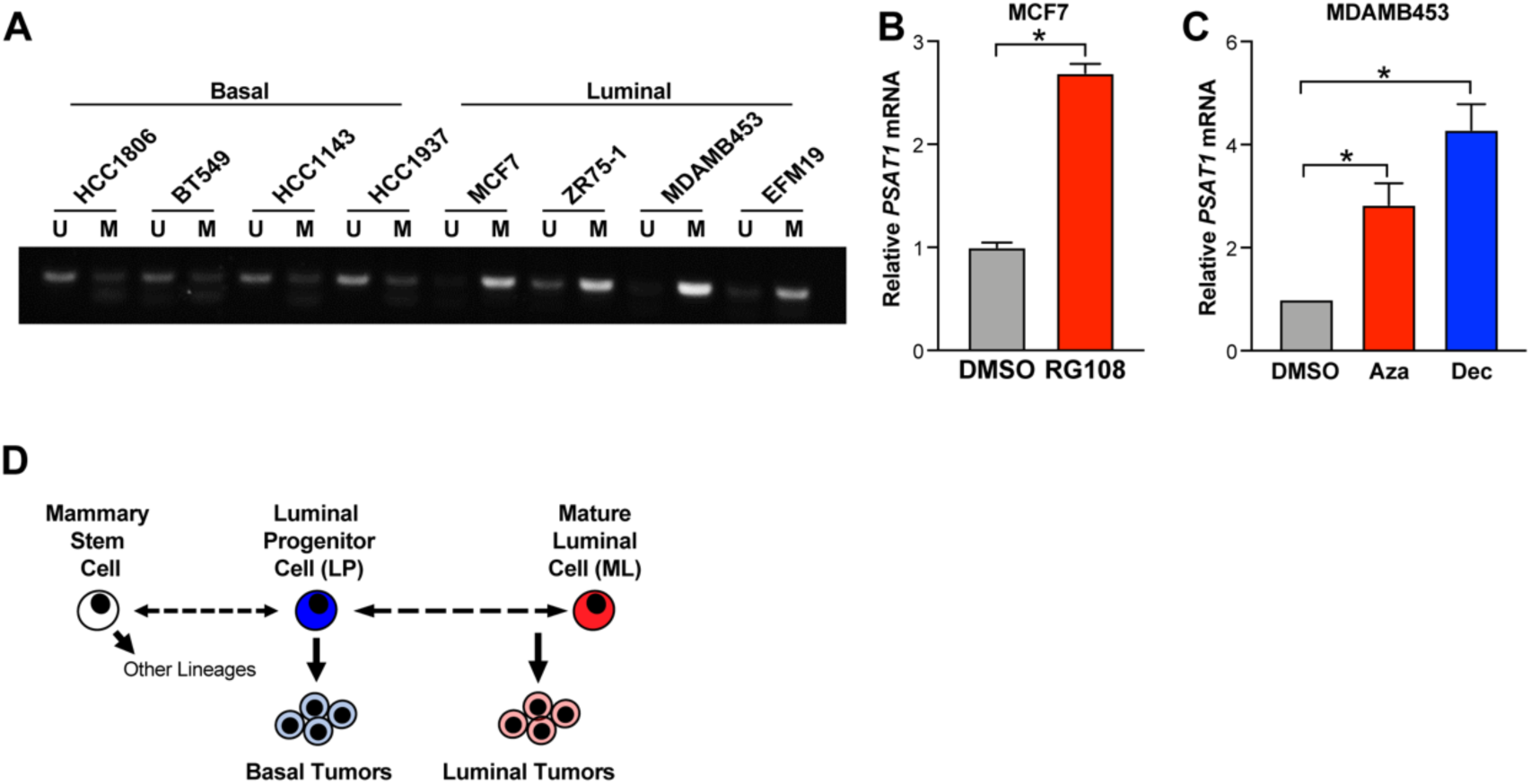
**Related to Figure 4.** **(A)** Representative methylation specific PCR detecting methylated (M) and unmethylated (U) *PSAT1* promoter DNA in basal and luminal breast cancer cells. **(B)** *PSAT1* mRNA in MCF7 cells treated with RG108 (10 µM for 3 days). Values are the mean +/- SD of triplicate samples from an experiment representative of three independent experiments. * indicates *p* < 0.05 in an unpaired two-sided t test. **(C)** PSAT1 mRNA level in MDAMB453 treated with azacytidine (5 µM for 3 days) or decitabine (1 µM for 3 days). Values are the mean ± SD of triplicate samples from an experiment representative of two independent experiments. * indicates *p* < 0.05 in an unpaired two-sided t test. **(D)** Simplified model of the mammary epithelial stem cell hierarchy.

## Notes

### Competing Interest Statement

The authors have declared no competing interest.

## REFERENCES

Andre, F., Ciruelos, E., Rubovszky, G., Campone, M., Loibl, S., Rugo, H.S., Iwata, H., Conte, P., Mayer, I.A., Kaufman, B., et al. (2019). Alpelisib for PIK3CA-Mutated, Hormone Receptor-Positive Advanced Breast Cancer. N Engl J Med 380, 1929–1940.

Barretina, J., Caponigro, G., Stransky, N., Venkatesan, K., Margolin, A.A., Kim, S., Wilson, C.J., Lehar, J., Kryukov, G.V., Sonkin, D., et al. (2012). The Cancer Cell Line Encyclopedia enables predictive modelling of anticancer drug sensitivity. Nature 483, 603–607.

Cancer Genome Atlas, N. (2012). Comprehensive molecular portraits of human breast tumours. Nature 490, 61–70.

Cantor, J.R., Abu-Remaileh, M., Kanarek, N., Freinkman, E., Gao, X., Louissaint, A., Jr., Lewis, C.A., and Sabatini, D.M. (2017). Physiologic Medium Rewires Cellular Metabolism and Reveals Uric Acid as an Endogenous Inhibitor of UMP Synthase. Cell 169, 258–272 e217.

Chen, J., Chung, F., Yang, G., Pu, M., Gao, H., Jiang, W., Yin, H., Capka, V., Kasibhatla, S., Laffitte, B., et al. (2013). Phosphoglycerate dehydrogenase is dispensable for breast tumor maintenance and growth. Oncotarget 4, 2502–2511.

Coloff, J.L., Murphy, J.P., Braun, C.R., Harris, I.S., Shelton, L.M., Kami, K., Gygi, S.P., Selfors, L.M., and Brugge, J.S. (2016). Differential Glutamate Metabolism in Proliferating and Quiescent Mammary Epithelial Cells. Cell Metab 23, 867–880.

Curtis, C., Shah, S.P., Chin, S.F., Turashvili, G., Rueda, O.M., Dunning, M.J., Speed, D., Lynch, A.G., Samarajiwa, S., Yuan, Y., et al. (2012). The genomic and transcriptomic architecture of 2,000 breast tumours reveals novel subgroups. Nature 486, 346–352.

DeNicola, G.M., Chen, P.H., Mullarky, E., Sudderth, J.A., Hu, Z., Wu, D., Tang, H., Xie, Y., Asara, J.M., Huffman, K.E., et al. (2015). NRF2 regulates serine biosynthesis in non-small cell lung cancer. Nat Genet 47, 1475–1481.

Diehl, F.F., Lewis, C.A., Fiske, B.P., and Vander Heiden, M.G. (2019). Cellular redox state constrains serine synthesis and nucleotide production to impact cell proliferation. Nat Metab 1, 861–867.

Finn, R.S., Martin, M., Rugo, H.S., Jones, S., Im, S.A., Gelmon, K., Harbeck, N., Lipatov, O.N., Walshe, J.M., Moulder, S., et al. (2016). Palbociclib and Letrozole in Advanced Breast Cancer. N Engl J Med 375, 1925–1936.

Gantner, M.L., Eade, K., Wallace, M., Handzlik, M.K., Fallon, R., Trombley, J., Bonelli, R., Giles, S., Harkins-Perry, S., Heeren, T.F.C., et al. (2019). Serine and Lipid Metabolism in Macular Disease and Peripheral Neuropathy. N Engl J Med 381, 1422–1433.

Gaude, E., and Frezza, C. (2016). Tissue-specific and convergent metabolic transformation of cancer correlates with metastatic potential and patient survival. Nat Commun 7, 13041.

Goetz, M.P., Toi, M., Campone, M., Sohn, J., Paluch-Shimon, S., Huober, J., Park, I.H., Tredan, O., Chen, S.C., Manso, L., et al. (2017). MONARCH 3: Abemaciclib As Initial Therapy for Advanced Breast Cancer. J Clin Oncol 35, 3638–3646.

Hoadley, K.A., Yau, C., Hinoue, T., Wolf, D.M., Lazar, A.J., Drill, E., Shen, R., Taylor, A.M., Cherniack, A.D., Thorsson, V., et al. (2018). Cell-of-Origin Patterns Dominate the Molecular Classification of 10,000 Tumors from 33 Types of Cancer. Cell 173, 291–304 e296.

Hortobagyi, G.N., Stemmer, S.M., Burris, H.A., Yap, Y.S., Sonke, G.S., Paluch-Shimon, S., Campone, M., Petrakova, K., Blackwell, K.L., Winer, E.P., et al. (2018). Updated results from MONALEESA-2, a phase III trial of first-line ribociclib plus letrozole versus placebo plus letrozole in hormone receptor-positive, HER2-negative advanced breast cancer. Annals of oncology : official journal of the European Society for Medical Oncology / ESMO 29, 1541–1547.

Hu, J., Locasale, J.W., Bielas, J.H., O’Sullivan, J., Sheahan, K., Cantley, L.C., Vander Heiden, M.G., and Vitkup, D. (2013). Heterogeneity of tumor-induced gene expression changes in the human metabolic network. Nat Biotechnol 31, 522–529.

Jiang, G., Zhang, S., Yazdanparast, A., Li, M., Pawar, A.V., Liu, Y., Inavolu, S.M., and Cheng, L. (2016). Comprehensive comparison of molecular portraits between cell lines and tumors in breast cancer. BMC Genomics 17 Suppl 7, 525.

Johansson, H.J., Socciarelli, F., Vacanti, N.M., Haugen, M.H., Zhu, Y., Siavelis, I., Fernandez-Woodbridge, A., Aure, M.R., Sennblad, B., Vesterlund, M., et al. (2019). Breast cancer quantitative proteome and proteogenomic landscape. Nat Commun 10, 1600.

Kim, M., and Costello, J. (2017). DNA methylation: an epigenetic mark of cellular memory. Exp Mol Med 49, e322.

Labuschagne, C.F., van den Broek, N.J., Mackay, G.M., Vousden, K.H., and Maddocks, O.D. (2014). Serine, but not glycine, supports one-carbon metabolism and proliferation of cancer cells. Cell reports 7, 1248–1258.

LeBoeuf, S.E., Wu, W.L., Karakousi, T.R., Karadal, B., Jackson, S.R., Davidson, S.M., Wong, K.K., Koralov, S.B., Sayin, V.I., and Papagiannakopoulos, T. (2020). Activation of Oxidative Stress Response in Cancer Generates a Druggable Dependency on Exogenous Non-essential Amino Acids. Cell Metab 31, 339–350 e334.

Lehmann, B.D., Bauer, J.A., Chen, X., Sanders, M.E., Chakravarthy, A.B., Shyr, Y., and Pietenpol, J.A. (2011). Identification of human triple-negative breast cancer subtypes and preclinical models for selection of targeted therapies. J Clin Invest 121, 2750–2767.

Li, L.C., and Dahiya, R. (2002). MethPrimer: designing primers for methylation PCRs. Bioinformatics 18, 1427–1431.

Lim, E., Vaillant, F., Wu, D., Forrest, N.C., Pal, B., Hart, A.H., Asselin-Labat, M.L., Gyorki, D.E., Ward, T., Partanen, A., et al. (2009). Aberrant luminal progenitors as the candidate target population for basal tumor development in BRCA1 mutation carriers. Nat Med 15, 907–913.

Locasale, J.W., Grassian, A.R., Melman, T., Lyssiotis, C.A., Mattaini, K.R., Bass, A.J., Heffron, G., Metallo, C.M., Muranen, T., Sharfi, H., et al. (2011). Phosphoglycerate dehydrogenase diverts glycolytic flux and contributes to oncogenesis. Nat Genet 43, 869–874.

Maddocks, O.D., Berkers, C.R., Mason, S.M., Zheng, L., Blyth, K., Gottlieb, E., and Vousden, K.H. (2013). Serine starvation induces stress and p53-dependent metabolic remodelling in cancer cells. Nature 493, 542–546.

Maddocks, O.D.K., Athineos, D., Cheung, E.C., Lee, P., Zhang, T., van den Broek, N.J.F., Mackay, G.M., Labuschagne, C.F., Gay, D., Kruiswijk, F., et al. (2017). Modulating the therapeutic response of tumours to dietary serine and glycine starvation. Nature 544, 372–376.

Mattaini, K.R., Sullivan, M.R., and Vander Heiden, M.G. (2016). The importance of serine metabolism in cancer. J Cell Biol 214, 249–257.

Mayers, J.R., Torrence, M.E., Danai, L.V., Papagiannakopoulos, T., Davidson, S.M., Bauer, M.R., Lau, A.N., Ji, B.W., Dixit, P.D., Hosios, A.M., et al. (2016). Tissue of origin dictates branched-chain amino acid metabolism in mutant Kras-driven cancers. Science 353, 1161–1165.

Mayers, J.R., and Vander Heiden, M.G. (2017). Nature and Nurture: What Determines Tumor Metabolic Phenotypes? Cancer Res 77, 3131–3134.

Méndez-Lucas, A., Lin, W., Driscoll, P.C., Legrave, N., Novellasdemunt, L., Xie, C., Charles, M., Wilson, Z., Jones, N.P., Rayport, S., et al. (2020). Identifying strategies to target the metabolic flexibility of tumours. Nat Metab 1, 335–350.

Molloy, M.E., White, B.E., Gherezghiher, T., Michalsen, B.T., Xiong, R., Patel, H., Zhao, H., Maximov, P.Y., Jordan, V.C., Thatcher, G.R., et al. (2014). Novel selective estrogen mimics for the treatment of tamoxifen-resistant breast cancer. Mol Cancer Ther 13, 2515–2526.

Muir, A., Danai, L.V., Gui, D.Y., Waingarten, C.Y., Lewis, C.A., and Vander Heiden, M.G. (2017). Environmental cystine drives glutamine anaplerosis and sensitizes cancer cells to glutaminase inhibition. Elife 6.

Mullarky, E., Lucki, N.C., Beheshti Zavareh, R., Anglin, J.L., Gomes, A.P., Nicolay, B.N., Wong, J.C., Christen, S., Takahashi, H., Singh, P.K., et al. (2016). Identification of a small molecule inhibitor of 3-phosphoglycerate dehydrogenase to target serine biosynthesis in cancers. Proc Natl Acad Sci U S A 113, 1778–1783.

Nilsson, L.M., Forshell, T.Z., Rimpi, S., Kreutzer, C., Pretsch, W., Bornkamm, G.W., and Nilsson, J.A. (2012). Mouse genetics suggests cell-context dependency for Myc-regulated metabolic enzymes during tumorigenesis. PLoS Genet 8, e1002573.

Nowak, M.A., Boerlijst, M.C., Cooke, J., and Smith, J.M. (1997). Evolution of genetic redundancy. Nature 388, 167–171.

Osborne, C.K., and Schiff, R. (2011). Mechanisms of endocrine resistance in breast cancer. Annu Rev Med 62, 233–247.

Pacold, M.E., Brimacombe, K.R., Chan, S.H., Rohde, J.M., Lewis, C.A., Swier, L.J., Possemato, R., Chen, W.W., Sullivan, L.B., Fiske, B.P., et al. (2016). A PHGDH inhibitor reveals coordination of serine synthesis and one-carbon unit fate. Nat Chem Biol 12, 452–458.

Pellacani, D., Bilenky, M., Kannan, N., Heravi-Moussavi, A., Knapp, D., Gakkhar, S., Moksa, M., Carles, A., Moore, R., Mungall, A.J., et al. (2016). Analysis of Normal Human Mammary Epigenomes Reveals Cell-Specific Active Enhancer States and Associated Transcription Factor Networks. Cell reports 17, 2060–2074.

Perou, C.M., Sorlie, T., Eisen, M.B., van de Rijn, M., Jeffrey, S.S., Rees, C.A., Pollack, J.R., Ross, D.T., Johnsen, H., Akslen, L.A., et al. (2000). Molecular portraits of human breast tumours. Nature 406, 747–752.

Possemato, R., Marks, K.M., Shaul, Y.D., Pacold, M.E., Kim, D., Birsoy, K., Sethumadhavan, S., Woo, H.K., Jang, H.G., Jha, A.K., et al. (2011). Functional genomics reveal that the serine synthesis pathway is essential in breast cancer. Nature 476, 346–350.

Rohde, J.M., Brimacombe, K.R., Liu, L., Pacold, M.E., Yasgar, A., Cheff, D.M., Lee, T.D., Rai, G., Baljinnyam, B., Li, Z., et al. (2018). Discovery and optimization of piperazine-1-thiourea-based human phosphoglycerate dehydrogenase inhibitors. Bioorg Med Chem 26, 1727–1739.

Selfors, L.M., Stover, D.G., Harris, I.S., Brugge, J.S., and Coloff, J.L. (2017). Identification of cancer genes that are independent of dominant proliferation and lineage programs. Proc Natl Acad Sci U S A 114, E11276–E11284.

Siegel, R.L., Miller, K.D., and Jemal, A. (2019). Cancer statistics, 2019. CA Cancer J Clin 69, 7–34.

Sorlie, T., Perou, C.M., Tibshirani, R., Aas, T., Geisler, S., Johnsen, H., Hastie, T., Eisen, M.B., van de Rijn, M., Jeffrey, S.S., et al. (2001). Gene expression patterns of breast carcinomas distinguish tumor subclasses with clinical implications. Proc Natl Acad Sci U S A 98, 10869–10874.

Sullivan, M.R., Mattaini, K.R., Dennstedt, E.A., Nguyen, A.A., Sivanand, S., Reilly, M.F., Meeth, K., Muir, A., Darnell, A.M., Bosenberg, M.W., et al. (2019). Increased Serine Synthesis Provides an Advantage for Tumors Arising in Tissues Where Serine Levels Are Limiting. Cell Metab.

Tardito, S., Oudin, A., Ahmed, S.U., Fack, F., Keunen, O., Zheng, L., Miletic, H., Sakariassen, P.O., Weinstock, A., Wagner, A., et al. (2015). Glutamine synthetase activity fuels nucleotide biosynthesis and supports growth of glutamine-restricted glioblastoma. Nat Cell Biol 17, 1556–1568.

Thiele, I., Swainston, N., Fleming, R.M., Hoppe, A., Sahoo, S., Aurich, M.K., Haraldsdottir, H., Mo, M.L., Rolfsson, O., Stobbe, M.D., et al. (2013). A community-driven global reconstruction of human metabolism. Nat Biotechnol 31, 419–425.

Vander Heiden, M.G., and DeBerardinis, R.J. (2017). Understanding the Intersections between Metabolism and Cancer Biology. Cell 168, 657–669.

Visvader, J.E., and Stingl, J. (2014). Mammary stem cells and the differentiation hierarchy: current status and perspectives. Genes Dev 28, 1143–1158.

Wang, Q., Liberti, M.V., Liu, P., Deng, X., Liu, Y., Locasale, J.W., and Lai, L. (2017). Rational Design of Selective Allosteric Inhibitors of PHGDH and Serine Synthesis with Anti-tumor Activity. Cell Chem Biol 24, 55–65.

Worton, K.S., Kerbel, R.S., and Andrulis, I.L. (1991). Hypomethylation and reactivation of the asparagine synthetase gene induced by L-asparaginase and ethyl methanesulfonate. Cancer Res 51, 985–989.

Yuneva, M.O., Fan, T.W., Allen, T.D., Higashi, R.M., Ferraris, D.V., Tsukamoto, T., Mates, J.M., Alonso, F.J., Wang, C., Seo, Y., et al. (2012). The metabolic profile of tumors depends on both the responsible genetic lesion and tissue type. Cell Metab 15, 157–170.

